# Meta16S: large-scale discovery and taxonomic assignment of unknown microbes from 16S amplicon sequencing samples

**DOI:** 10.64898/2026.05.19.726236

**Authors:** Fabio Cumbo, Giovanni Felici, Daniel Blankenberg, Federica Valeriani, Vincenzo Romano Spica, Daniele Santoni

## Abstract

**Background:** The exponential growth of public metagenomic datasets offers an unprecedented opportunity to explore microbial diversity. However, analyzing this vast amount of data presents significant computational challenges. While shotgun metagenomics provides deep functional and taxonomic resolution, its high cost still limits its application. On the other hand, 16S rRNA gene sequencing remains a cost-effective and widely used alternative, but tools are needed to maximize its discovery potential. Traditional clustering is not scalable, obstructing the creation of a comprehensive and continuously updated catalog of microbial life from 16S data.

**Methods:** We developed a reproducible and scalable Snakemake pipeline for the incremental clustering of 16S rRNA amplicons. The workflow begins by constructing a reference database from bacterial and archaeal genomes. It then processes 16S rRNA samples sequentially. For each new sample, sequences are first mapped against the existing cluster centroids. Sequences that match known centroids are assigned accordingly, while unmapped sequences are clustered independently to form novel operational taxonomic units (OTUs). These new centroids are then merged with the existing database, allowing it to grow dynamically without the need for computationally prohibitive all-at-once re-clustering.

**Results:** Our pipeline enables the efficient and continuous expansion of a 16S rRNA cluster database. By processing a large corpus of public 16S rRNA samples, we generated a comprehensive atlas of tens of thousands of OTUs. A significant fraction of these clusters, particularly at the genus and family levels, were classified as unknown.

**Conclusions:** This work provides a powerful, open-source tool for large-scale analysis of 16S rRNA samples. The incremental clustering strategy overcomes the scalability limitations of traditional methods, allowing researchers to leverage public data and discover novel microbes in their own microbiome samples.

## INTRODUCTION

For decades, the 16S ribosomal RNA (rRNA) gene has served as a cornerstone of microbial ecology. Its central role dates back to the 1970s, when Woese and Fox published their seminal work [1], paving the way for molecular, sequence-based taxonomic classification, later formalized and expanded in a subsequent study [2,3]. The 16S rRNA mosaic of conserved and hypervariable regions makes it an ideal molecular marker for taxonomic classification, allowing researchers to survey the composition of complex microbial communities from nearly any environment [4–7]. From the human gut to the deep-sea vents, 16S rRNA gene sequencing has provided fundamental insights into the structure, function, and dynamics of the microbial world, forming the bedrock of countless ecological and clinical studies [3,8,9].

The 16S rRNA gene contains nine distinct hypervariable regions (V1–V9), and the choice of which region to amplify heavily influences the taxonomic resolution of a study [5,7,10–12]. While regions like V3–V4 are frequently targeted in broad microbiome surveys, recent evaluations have highlighted the superior discriminatory power of other segments depending on the sample type and targeted taxa. For instance, López-Aladid et al. (2023) [10] demonstrated that the V1–V2 hypervariable region offers the most accurate taxonomic identification for respiratory samples, underscoring its robustness for deep, species-level microbial profiling. Furthermore, the V1–V2 region has proven highly effective in distinguishing closely related species across diverse environmental and clinical contexts, often outperforming other regions in its ability to resolve complex phylogenetic relationships [5,7].

To interpret these amplicon sequences, modern bioinformatics pipelines such as DADA2 [13], QIIME 2 [14], and mothur [15] have become standard tools in the field. They provide end-to-end frameworks for sequence processing, chimera removal, and taxonomic assignment. Notably, tools like DADA2 explicitly create Amplicon Sequence Variants (ASVs) via sequence denoising, which can be directly compared across studies and later optionally clustered into operational taxonomic units (OTUs). These pipelines rely heavily on curated reference databases, including SILVA [16], Greengenes [17], and the Ribosomal Database Project (RDP) [18]. Together, they have become standard tools in microbial ecology and microbiome research, enabling reproducible and scalable analyses across diverse environments.

We are currently living in an unprecedented era of biological data generation. The proliferation of high-throughput sequencing technologies has led to an exponential increase in the volume of publicly available data, particularly within metagenomics. Public repositories like the Sequence Read Archive (SRA) [19–21] of the National Institutes of Health (NIH) now host petabytes of sequencing information, representing a vast and largely untapped resource for biological discovery [22]. This digital bank offers a remarkable opportunity to build a truly comprehensive census of microbial life on Earth, moving beyond the limitations of individual studies to synthesize knowledge on a global scale.

However, harnessing this potential requires navigating a critical trade-off between analytical depth and practical accessibility. Shotgun metagenomics, which sequences the entire genomic content of a community, offers unparalleled resolution, providing insights into both taxonomic composition and functional potential [23–27]. Yet, its relatively high cost remains a significant barrier, often placing it beyond the reach of smaller laboratories or large-scale screening projects with limited budgets. In this context, 16S rRNA gene sequencing still represents a powerful and relevant technique. Its cost-effectiveness, established protocols, and lower computational requirements make it a pragmatic choice for a broad spectrum of research. It effectively democratizes discovery, enabling a wider scientific community to participate in the exploration of microbial ecosystems.

Despite this wealth of available data, a significant limitation of standard reference-based classification is its inability to characterize “microbial dark matter”, the vast portion of the microbial world that remains uncultivated and absent from reference databases [28–35]. Previous large-scale environmental surveys have reportedly revealed that immense swaths of phylogenetic diversity remain unmapped, with unknown microbes dominating many environmental niches and playing crucial roles in biogeochemical cycles [36]. When relying solely on static, pre-computed reference databases, sequences originating from these unknown organisms are either discarded as unclassified or forced into incorrect taxonomic bins. This obscures true community structures and severely hinders the discovery of novel taxa from accumulating public data.

Furthermore, large-scale *de novo* analysis of 16S data is affected by different computational bottlenecks. Traditional methods for designing Operational Taxonomic Units (OTUs – i.e., clusters of similar sequences that serve as proxies for microbial species) rely on algorithms that require all sequences to be processed simultaneously. Tools like UCLUST [37] or CD-HIT [38] perform an all-vs-all comparison, a task whose computational demand grows quadratically with the number of sequences. This approach is fundamentally unscalable. Adding even a single new sample necessitates a complete re-clustering of the entire dataset, a process that quickly becomes computationally infeasible as databases grow from thousands to millions of samples. This bottleneck has severely limited our ability to create a single, unified, and continuously updated reference catalog of microbial diversity.

To overcome this challenge, we propose a paradigm shift from static, all-at-once processing to a dynamic, incremental clustering strategy. The core concept is to build a reference database that grows organically as new data becomes available, without the need to re-analyze previously processed samples. Rather than performing a quadratic all-vs-all comparison, our approach maps new sequences against an existing centroid database, a process that scales proportionally to the number of incoming reads *N* and the current number of centroids *K* (*O*(*N* × *K*)). Only the unmapped fraction, representing potential novel diversity, undergoes *de novo* clustering. Thus, the overall computational demand scales sub-quadratically, allowing the pipeline to seamlessly process thousands of samples and continuously expand the catalog of known life.

In this paper, we present Meta16S, a novel, fully reproducible, and scalable Snakemake [39] workflow designed to address these challenges. Our contribution is threefold. First, we have developed a scalable pipeline capable of incrementally clustering an ongoing influx of 16S rRNA amplicon samples without prohibitive computational overhead. Second, we utilized this pipeline to construct a comprehensive and dynamically growing cluster database that systematically catalogs both known taxonomic groups and uncharacterized “microbial dark matter”. By applying this framework to a large corpus of publicly available 16S rRNA data spanning human gut, oral, and environmental water ecosystems, we showcase the profound extent of novel microbial diversity uncovered. We further demonstrate how these newly discovered, unannotated OTUs exhibit distinct environmental enrichment patterns, highlighting their ecological significance and proving the value of a continuously evolving global census of microbial life.

In the following sections, we will first detail the architecture of our Snakemake pipeline and the incremental clustering methodology. We will then present the results of applying our workflow to a large public dataset, quantifying the growth of the cluster database and analyzing the structural distribution of known and novel microbial diversity. Finally, we will discuss the implications of these findings for future microbiome research.

## MATERIALS AND METHODS

This section details the computational framework developed for the incremental clustering of 16S rRNA sequences and the subsequent integration into quantitative profiling tools. We describe the overall architecture of our pipeline, managed by Snakemake, and the specific roles of the key bioinformatics software employed. We then outline the step-by-step logic of the workflow, from the initial construction of a reference database to the iterative process of mapping, clustering, and merging that allows the database to expand.

### Dataset

To evaluate the performance of the Meta16S pipeline and build our dynamic cluster database, a total of 734 16S rRNA samples were analyzed in this study. The dataset was curated to encompass a diverse range of microbial niches, allowing us to assess environmental enrichment patterns. The vast majority of the samples originated from the human gut microbiome (*n* = 691), complemented by additional subsets representing oral (*n* = 11) and environmental water (*n* = 32) ecosystems.

Furthermore, to ensure the robustness of our framework and assess the potential impact of sequencing technology variability on the clustering process, the samples were sourced from two distinct Illumina platforms. The bulk of the sequencing data was generated using the Illumina iSeq platform (*n* = 718). A smaller control group consisting of samples sequenced on the Illumina MiSeq platform (*n* = 16) was specifically included to determine whether platform-specific biases or error profiles affect the *de novo* discovery and taxonomic assignment workflow.

### Workflow Architecture

The entire analytical process presented in this study is encapsulated into a robust and portable computational workflow, designed for scalability, reproducibility, and accessibility to researchers with varying levels of computational expertise.

The core of our workflow is orchestrated using Snakemake, a powerful and flexible workflow management system. Snakemake formalizes the entire analysis as a series of rules, where each rule defines how to produce output files from input files by running a specific piece of software. This rule-based structure provides several key advantages. First, it ensures reproducibility by explicitly defining every step, parameter, and dependency. Second, it enables scalability, as Snakemake can automatically parallelize independent tasks across multiple CPU cores or submit them to high-performance computing (HPC) clusters. Finally, it offers efficiency: if a parameter is changed, Snakemake is intelligent enough to re-run only the affected downstream steps, saving significant computational time. The workflow is defined in a Snakefile, which serves as the central, executable blueprint for the entire analysis.

To guarantee a consistent and reproducible software environment across different operating systems and computational infrastructures, all software dependencies are managed through the Conda package manager [40]. A comprehensive environment.yaml file is provided with the workflow, which lists the exact versions of all required tools, including VSEARCH (v2.30.0) [41], barrnap (v0.9), and ncbitax2lin (v2.4.1). When the pipeline is executed, Snakemake automatically creates an isolated Conda environment from this file, ensuring that the analysis runs with the precise software stack used in this study, thereby eliminating machine-dependent issues.

The workflow architecture is summarized in Figure 1 below.

**Figure 1:**
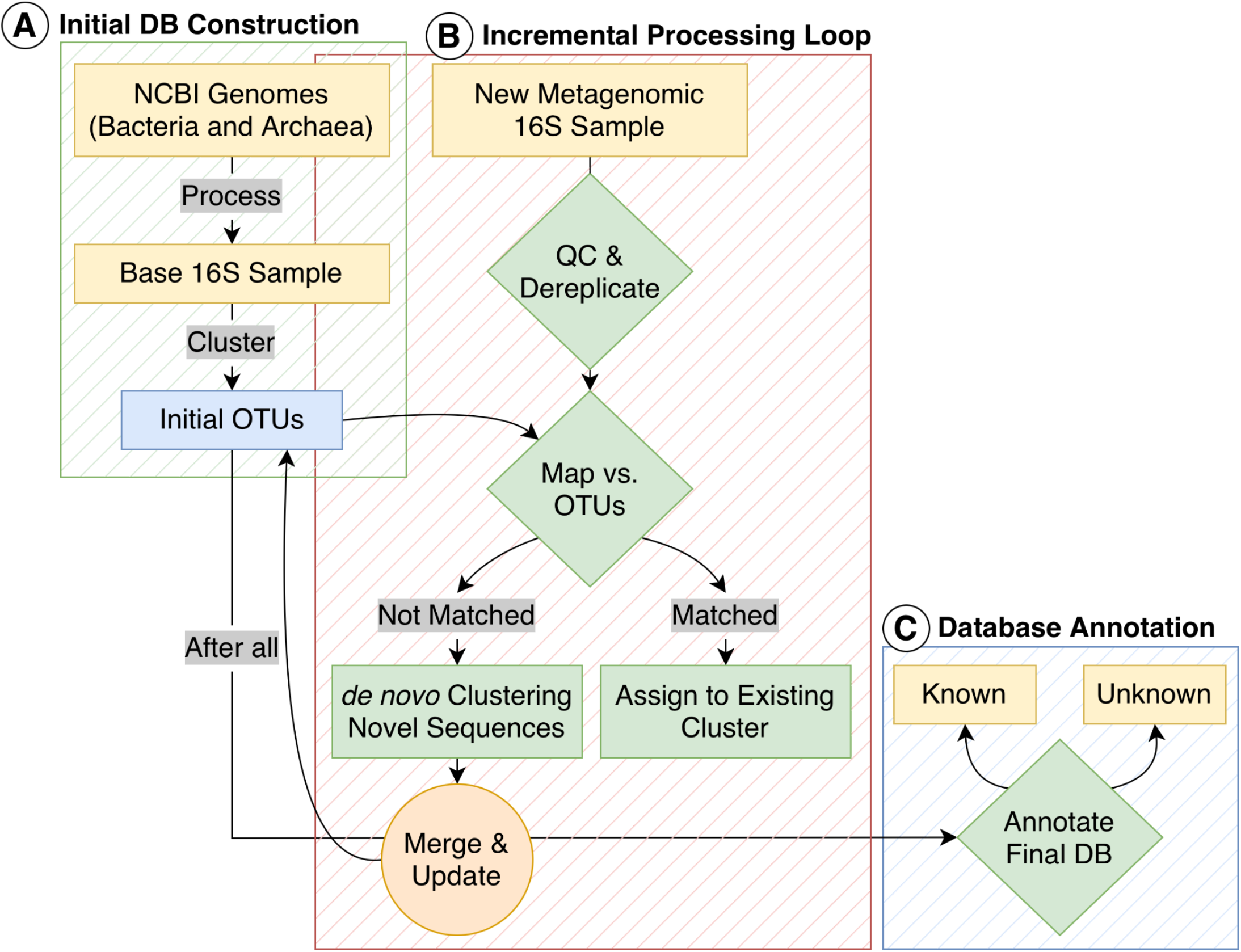
Schematic overview of the incremental clustering and quantitative profiling workflow. This diagram illustrates the complete data flow of the pipeline. <u>A – Initial DB Construction:</u> reference genomes from NCBI GenBank are processed with barrnap and seqkit to create a base artificial sample of 16S sequences that are clustered to form the initial version of the OTU database; <u>B – Incremental Processing Loop:</u> for each new 16S rRNA sample, reads are quality-filtered and dereplicated. They are then mapped against the current OTU database. Matched reads are assigned to existing clusters, while non-matching reads are clustered *de novo* to form new OTUs. The centroids of these new OTUs are merged with the main database, causing it to expand. This loop is repeated for all subsequent samples; <u>C – Database Annotation:</u> once all samples are processed, the final, comprehensive OTU database is annotated. Clusters containing reference sequences are labeled “known” with a consensus taxonomy, while clusters without references are labeled “unknown”.

### Construction of the Initial Reference Database

The foundation of our incremental clustering strategy is a robust, high-quality reference database containing known 16S rRNA gene sequences. This database serves two critical purposes: it provides the initial set of centroids for clustering and acts as the taxonomic anchor for annotating newly formed clusters. The construction of this database was performed as follows:

1. <u>Source genome retrieval:</u> a comprehensive set of prokaryotic genomes was downloaded from the NCBI GenBank repository. This retrieval was automated using the *get_ncbi_genomes*.*py* subroutine as part of the MetaSBT framework [42]. This script was configured to retrieve one representative genome per species across both the Bacteria and Archaea microbial domains. Crucially, the subroutine extends the classic definition of a reference genome by implementing a more nuanced selection logic. To select the best possible representative for each species, it evaluates genomes based on specific RefSeq exclusion criteria. This ensures that even for species lacking a formal reference tag, an assembly is considered if it passes the selection criteria. The specific selection criteria used by the subroutine are detailed in Table 1;
2. <u>16S rRNA genome prediction and extraction:</u> since the downloaded files contain complete genome sequences, it was necessary to first identify and extract the 16S rRNA gene marker. This was accomplished using barrnap, a specialized tool for rapid rRNA gene prediction in bacterial and archaeal genomes. For each genome, barrnap was run with the appropriate kingdom flag (*--kingdom bac* or *--kingdom arc*) to optimize the accuracy of its underlying models, generating a FASTA file containing the full-length 16S rRNA gene sequences for that organism;
3. <u>Hypervariable region isolation:</u> the experimental data targeted in this workflow consists of amplicons from the V1-V2 hypervariable regions of the 16S rRNA gene. To ensure that our reference sequences were directly comparable to the experimental sequences, we isolated this specific region from the full-length reference genes. This was performed using the seqkit toolkit [43]. The *seqkit subseq* command was used to extract the nucleotide sequence spanning the approximate coordinates of the V1-V2 region (positions 60-250), effectively trimming the flanking conserved regions;
4. <u>Formatting of the base artificial sample:</u> all extracted V1-V2 reference sequences from every genome were concatenated into a single FASTQ file. To maintain compatibility with the rest of the workflow, which expects FASTQ input, a uniform, high-quality Phred score (ASCII character I, corresponding to a score of 40) was assigned to all bases. This consolidated file was designed as the base sample and serves as the essential seed input for the initial clustering step of the pipeline, forming the first version of the dynamic OTU database.

**Table 1:** Hierarchical criteria for the selection of reference genomes. This table outlines the selection logic implemented by the *get_ncbi_genomes*.*py* subroutine of MetaSBT to retrieve the single best representative genome per species from NCBI GenBank. Assemblies are filtered to exclude those with specific RefSeq categories that may indicate potential quality issues, such as being derived from a single cell or a metagenome. This process extends the formal definition of reference genome.

| RefSeq exclusion annotation | Long description |
| --- | --- |
| *derived from single cell* | This genome assembly was generated from material amplified from a single cell. Due to the nature of single-cell sequencing, the accuracy of the genome sequence may be affected. |
| *assembly from type material* | The genome assembly was constructed using sequences obtained from designated type material. |
| *assembly from synonym type material* | Analogously to *assembly from type material*, the genome assembly was constructed with sequences from designated synonym type material. |
| *assembly designated as neotype* | Sequences for the assembly were derived from neotype material. |
| *assembly designated as reftype* | Reftype material was used to derive sequences for constructing the genome assembly. |
| *assembly from pathotype material* | Here, the genome assembly was constructed with sequences from pathotype material. |
| *assembly from proxytype material* | Genome assembly was constructed using sequences from proxytype material. |
| *missing strain identifier* | No information about the strain and isolate of this genome assembly has been specified. |
| *genus undefined* | The absence of a genus designation in the lineage information makes it impossible to determine the precise taxonomic placement. |
| *from large multi-isolate project* | This assembly is part of a larger project that has produced over 100 assemblies, representing multiple isolates of the same species. |

### Per-Sample Processing of 16S rRNA Sequences

Prior to clustering, each individual 16S rRNA sample underwent a standardized pre-processing pipeline to ensure data quality and to format sequences for downstream analysis. The pipeline consists of three main steps: quality filtering, dereplication, and chimera removal.

1. <u>Quality filtering and read renaming:</u> raw forward reads (R1) from each sample were subjected to stringent quality control using the *vsearch --fastq_filter* command. Reads were discarded if they had a maximum number of expected errors (*--fastq_maxee*) greater than 1.0, a length shorter than 150 bp (*--fastq_minlen*) or longer than 300 bp (*--fastq_maxlen*), or if they contained any ambiguous bases (N, *--fastq_maxns 0*). This step is critical for removing low-quality and erroneous sequences that could otherwise lead to the formation of spurious OTUs. Immediately following filtering, the sequence headers were modified to prepend the sample name (e.g., *>sample1_read1*). This ensures that every read across the entire dataset has a unique identifier that traces back to its sample of origin, which is essential for downstream tracking and analysis;
2. <u>Dereplication:</u> the filtered, high-quality reads were dereplicated using the *vsearch --derep_fulllength* command. This process identifies and collapses all identical sequences into a single, unique representative sequence. The header of this representative sequence is then annotated with a *;size=N* tag, where *N* is the total number of identical reads found in that sample. Dereplication significantly reduces computational load for subsequent steps by removing redundancy while preserving quantitative information;
3. <u>De novo chimera removal:</u> chimera sequences, which are artifacts of PCR amplification where two or more distinct parent sequences are incorrectly joined, are a common source of inflated diversity estimates. To address this, we performed *de novo* chimera detection on the set of unique, dereplicated sequences using the *vsearch --uchime_denovo* command. This algorithm identifies chimeras by checking if a sequence appears to be a hybrid of two more abundant parent sequences within the same sample. All sequences flagged as chimeric were discarded, resulting in a final, clean set of non-chimeric sequences for each sample that were carried forward to the incremental clustering stage.

### The Incremental Clustering Algorithm

The central step of this workflow is an incremental clustering algorithm designed to expand the OTU database with each new sample, thereby avoiding the computationally prohibitive need to re-cluster the entire dataset. This process is executed as a loop, beginning with an initial seeding step followed by an iterative expansion for every subsequent sample. The identity threshold for all clustering and mapping steps was set at 95%, a commonly used proxy for genus-level OTUs.

1. <u>Initial database seeding:</u> the process is initiated using the base artificial sample, which contains the V1-V2 sequences from all reference genomes. This file is clustered using the standard *vsearch --cluster_fast* command. The resulting centroids from this initial step form version 1.0 of our dynamic OTU database, representing the known diversity derived from reference genomes;
2. <u>Iterative expansion loop:</u> for every subsequent 16S rRNA sample, the following three-part process is executed to integrate its sequences into the database (as illustrated in Figure 2):
  a. <u>Mapping against existing centroids:</u> first, the clean, non-chimeric sequences from the new sample are mapped against the most current version of the OTU centroid database using the rapid *vsearch --usearch_global* command. This step efficiently identifies and assigns any sequence that matches an existing centroid within the 95% identity threshold. Reads that are successfully mapped are considered part of known OTUs. The crucial output of this step is the set of reads that do not match any existing centroid;
  b. <u>*De novo* clustering of novel sequences:</u> the non-matching sequences, representing potential novel diversity, are passed to a separate, smaller clustering step. Using the *vsearch --cluster_size* command, these sequences are clustered among themselves. This targeted *de novo* clustering is computationally inexpensive as it operates only on a small fraction of the total data. The centroids generated from this step represent the novel OTUs discovered exclusively within the current sample;
  c. <u>Database merging and update:</u> finally, the main OTU database is updated. The centroids of the newly discovered OTUs (from step 2.b) are concatenated with the centroids from the previous version of the database. This combined FASTA file is then subjected to a final dereplication step (*vsearch --derep_fulllength*). This ensures that if a new centroid is identical to an existing one (a rare but possible edge case), it is merged, creating a single, non-redundant, and comprehensive OTU database. This updated database then serves as the reference for the mapping step 2.a of the next sample in the queue, thus completing the incremental loop.

**Figure 2:**
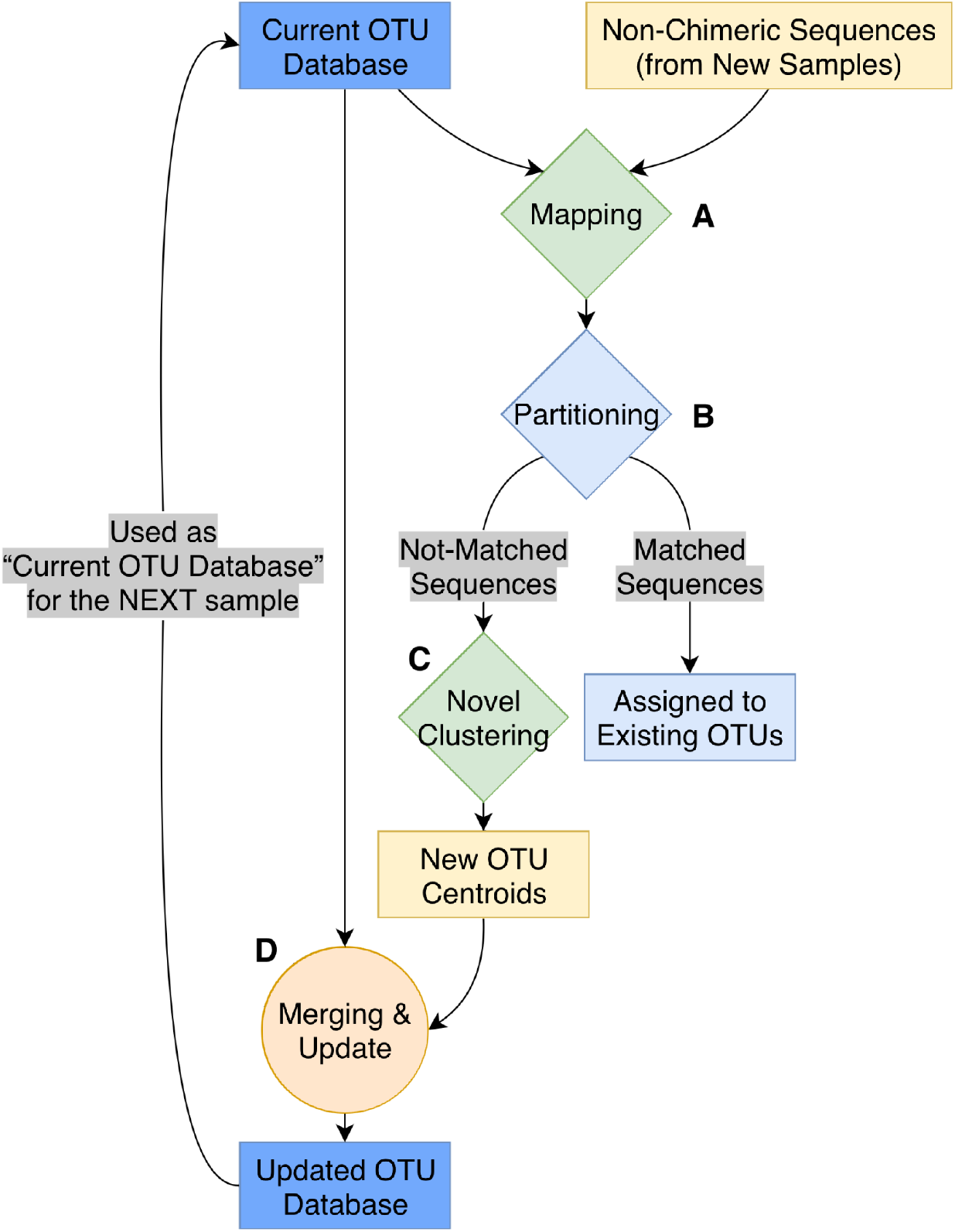
The incremental clustering loop. This flowchart details the iterative process for expanding the OTU database with each new sample. <u>A – Mapping:</u> non-chimeric sequences from a new sample are mapped against the current OTU database; <u>B – Partitioning:</u> sequences are partitioned into two groups: those that matched existing centroids and those that did not; <u>C – Novel clustering:</u> the non-matched sequences are clustered *de novo* to identify new OTUs; <u>D – Merging:</u> the centroids of these new OTUs are merged with the previous database and dereplicated to create the updated OTU database, which is then used as the reference for the next sample. This cycle of mapping, clustering novel sequences, and merging allows the database to grow dynamically without re-analyzing all prior data.

### Taxonomic Assignment and Annotation

After all samples were processed through the incremental clustering pipeline, a final annotation step was performed to assign a taxonomic identity to each OTU cluster and to formally classify it as known or unknown. This process integrates all clustering information generated throughout the workflow to create a final, unified summary.

1. <u>Aggregation of clustering evidence:</u> all mapping and clustering output files (in .*uc* format) from every step of the pipeline, including the initial base clustering, all incremental mapping steps, and all novel clustering steps, were concatenated into a single file. This file represents the complete history of all sequence-to-centroid assignments across the entire dataset;
2. <u>Creation of a definitive sequence-to-cluster map:</u> from the aggregated .*uc* file, a definitive map was constructed to link every unique sequence identifier to a single, final cluster ID. Since a sequence could appear multiple times in the raw output (e.g., as a hit in one context and a centroid in another), a simple rule was applied: the first assignment encountered for any given sequence was considered its definitive one. This created an unambiguous two-column table mapping every processed sequence to its final parent OTU;
3. <u>Taxonomic annotation via reference sequences:</u> the taxonomic identity of each OTU cluster was determined exclusively by the reference genome sequences it contained. For each cluster, we identified all member sequences that originated from our initial reference database. To assign a consensus taxonomy for the entire OTU, we implemented a strict 75% majority-voting threshold. Specifically, a taxonomic lineage was assigned to a given cluster if and only if at least 75% of the reference sequences within that cluster shared that exact classification. This lineage was truncated to the genus-level, providing a clear and consistent annotation level;
4. <u>Novelty classification:</u> based on the presence and absence of reference sequences, each OTU was assigned a final classification:
  a. **Known**: a cluster was labeled “known” if it contains at least one sequence derived from the initial reference genome database. The assigned taxonomy for these clusters is therefore anchored to a known organism;
  b. **Unknown**: a cluster was labeled “unknown” if it contained zero sequences from the reference database and was composed entirely of sequences from the processed samples. These clusters represent putative novel taxa that have no close relatives (at the 95% identity level) in the comprehensive reference genome collection, representing the primary targets of discovery for this workflow.

## RESULTS

Here, we present the outcomes of applying Meta16S to construct a comprehensive atlas of microbial diversity. We begin by characterizing the foundational reference database, detailing the extraction and processing of 16S rRNA V1-V2 regions from over 50 thousand prokaryotic genomes. We then describe the initial clustering of these reference sequences, which established the backbone of known diversity within our system. Subsequently, we demonstrate the pipeline’s capacity for discovery by processing a large corpus of 16S rRNA samples, resulting in a significant expansion of the cluster database with new, previously unannotated OTUs. Finally, we analyze the structure and distribution of this expanded diversity through principal component analysis (PCA) and examine the unknown cluster enrichment within the analyzed samples.

### From reference genomes to 16S rRNA and hypervariable regions: genera, genomes and 16S rRNA genes

A total of 66,360 representative genomes, consisting of one genome per species, were downloaded from NCBI GenBank, encompassing both the Bacteria (98.21%) and Archaea (1.79%) domains. To identify the 16S rRNA genes within these genomic assemblies, we utilized barrnap (Bacterial ribosomal RNA predictor). Given that sequence integrity is essential for precise clustering and subsequent taxonomic classification, the barrnap output was subjected to rigorous filtering to eliminate any sequence labeled as “partial”. This step was implemented to exclude truncated sequences, which frequently manifest at the margins of genomic contigs and risk generating substantial alignment artifacts, thereby yielding exclusively full-length, intact 16S rRNA gene predictions.

Subsequent to this filtering step, the V1-V2 hypervariable region was extracted from the complete 16S rRNA genes. This was achieved by targeting slightly broadening coordinates spanning the V1-V2 segment with each complete sequence, approximately corresponding to positions 27 through 338 according to the canonical *Escherichia coli* 16S rRNA numbering system. These wider boundaries were intentionally selected over the strict 60-250 range to guarantee that the complete hypervariable region was successfully captured across all diverse taxa. Because the downstream clustering process relies on sequence alignment, the inclusion of these short, conserved flanking bases does not negatively impact the final cluster delineation.

Ultimately, this procedure yielded 126,933 high-quality 16S rRNA gene predictions and their corresponding V1-V2 segments from the initial set of representative genomes. As an incidental finding beyond the primary scope of this study, barrnap identified an average of more than two 16S rRNA genes per genome, with certain genomes harboring dozens of identical copies. Moreover, multiple 16S rRNA genes housed within a single genome were typically identical or shared exceptionally high sequence similarity. As elaborated in the following section, fewer than 3% of the analyzed genomes contained V1-V2 sequences with sufficient divergence to be partitioned into distinct 95% similarity clusters. To mitigate any potential overrepresentation bias during our downstream cluster annotation, a dereplication strategy was applied: only a single representative V1-V2 16S sequence was retained per genome whenever multiple copies were assigned to the same cluster. This refinement process generated a final, curated reference dataset comprising 59,390 V1-V2 rRNA sequence distributed across 4,046 unique genera, as defined by the NCBI taxonomy.

### Clustering of 16S V1-V2 reference sequences and annotation

As detailed in the Materials and Methods section, VSEARCH was employed to cluster sequences derived from the NCBI reference genomes. The sequence identity threshold for cluster delineation was established at 95%, in accordance with benchmarks established by previous studies [6,44–47].

This clustering process yielded a total of 12,150 clusters. Of these, 10,652 clusters were homogeneous at the genus level, containing sequences derived exclusively from a single genus. Within this homogeneous subset, 6,981 were trivially singletons, 1,625 consisted of exactly two sequences from the same genus, and the remaining 2,046 clusters contained between 3 and 287 sequences assigned to the same genus.

An additional 500 clusters were highly dominated by a single genus, with over 75% of their constituent sequences sharing the same generic assignment. Consequently, a total of 11,152 clusters (representing 91.8% of the dataset, calculated as the sum of the 10,652 homogeneous clusters and the 500 genus-dominated clusters) can be unambiguously associated with a specific genus, as a minimum of three out of four, on average, of their sequences share the same taxonomic origin. Collectively, these 11,152 confidently assigned clusters encompass 46,907 sequences (accounting for 78.98% of the total 59,390 sequences analyzed). Within these specific clusters, 45,610 sequences (representing 76.79% of the entire dataset) correspond precisely to the dominant genus (i.e., the genus exhibiting a frequency greater than 75% within its respective cluster).

Figure 3 shows the relationship between the number of clusters and the number of genera they map to, based on the 75% assignment criterion. Specifically, the x-axis represents the number of clusters associated with a genus, while the y-axis indicates how many genera are associated with that number of clusters. Interestingly the plot in Figure 3 shows typical slope in the distribution of number of clusters per number of genera.

**Figure 3:**
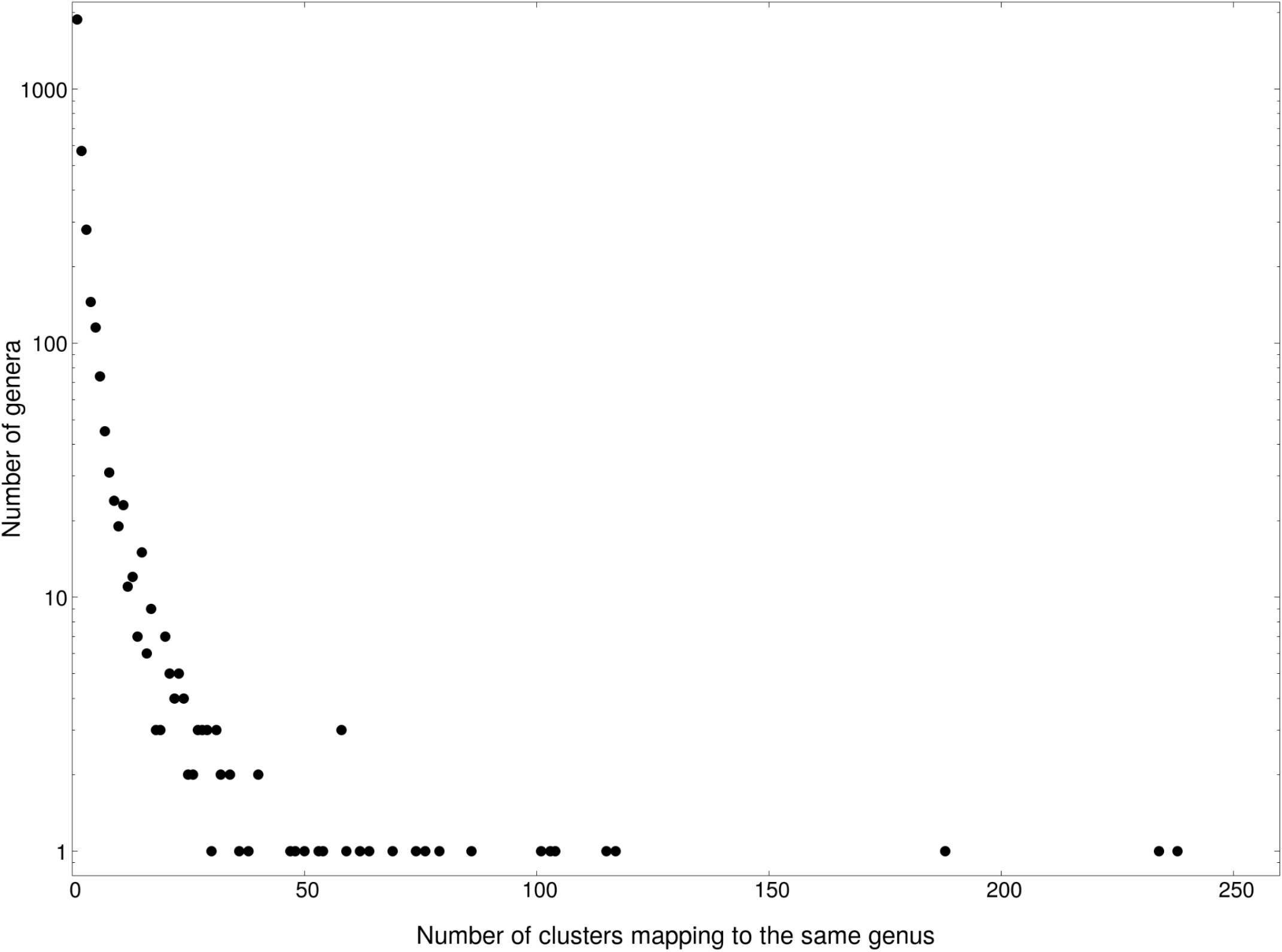
Distribution of the number of clusters associated with each genus (75% criterion). The x-axis represents the number of clusters per genus, while the y-axis indicates how many genera exhibit that association.

The remaining 998 clusters comprise 12,483 sequences (21.02% of the total 59,390). These clusters consist of sequences originating from multiple genera without a clear dominant majority, as the frequency of the most prevalent genus in these groups remains below the 75% threshold.

### Statistics on unknown clusters

Following the annotation of clusters derived from the 16S rRNA V1-V2 regions of reference genomes, the database was expanded by integrating sequences from environmental samples, in accordance with the methodology outlined in the preceding section.

During this phase, reads were either assigned to pre-existing, reference-annotated clusters or organized into newly generated clusters utilizing an incremental clustering approach. In total, we processed 734 samples, comprising an aggregate of 3,952,819 reads.

This integration yielded a final comprehensive dataset of 43,749 clusters. As previously noted, 12,150 of these clusters contained at least one known reference sequence. The size distribution for both the annotated (blue) and unknown (red) clusters is shown on a log-log scale in Figure 4. The figure illustrates the total number of clusters (y-axis) as a function of their respective size (x-axis). Notably, this distribution demonstrates a characteristic power-law decay, a trend emphasized by two black lines overlaid on the log-log representation.

**Figure 4:**
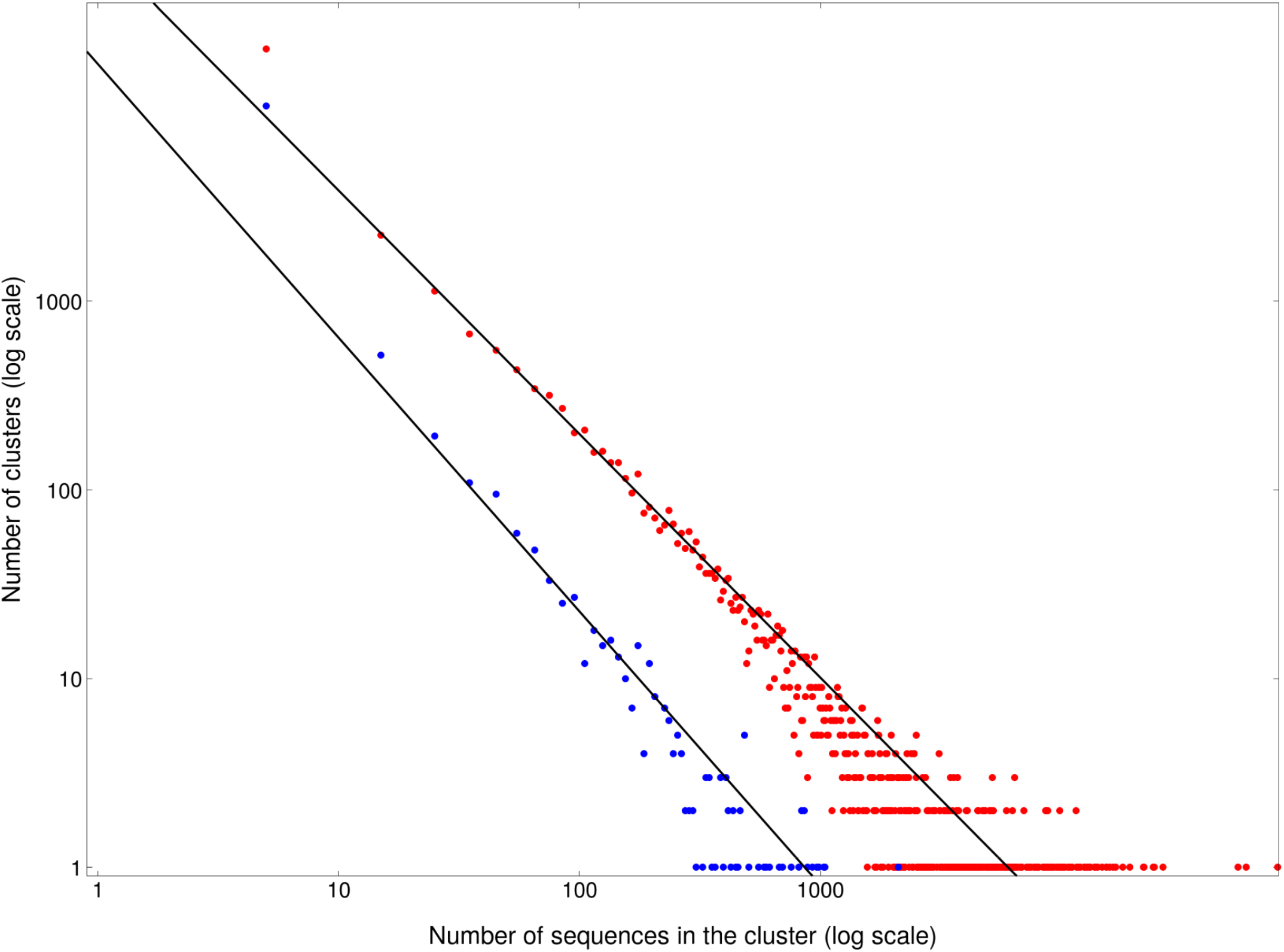
Cluster size distribution. The y-axis shows the number of clusters, while the x-axis represents the cluster size. The log-log plot highlights a characteristic power-law decay, emphasized by the overlaid black lines.

### Principal Component Analysis of clusters (based on centroids)

To visualize the distribution of both annotated and unknown clusters within the 16S rRNA V1-V2 sequence space, each cluster was represented by its centroid, defined as the sequence exhibiting the lowest average distance to all other sequences within that cluster. These centroids were subsequently projected onto a two-dimensional plane using Principal Component Analysis (PCA) derived from pairwise alignment distances. Given that computing all possible pairwise distances across the entire dataset was computationally prohibitive, a representative subsample of 2,000 clusters was randomly selected. Pairwise alignment distances were calculated exclusively for the centroids of this subsample. Consecutively, each cluster centroid was embedded in a distance space comprising 2,000 observations (the clusters/centroids) and 2,000 variables (their respective distances to all other sampled clusters/centroids). The PCA was then executed and visualized using the prcomp, ggbiplot, and ggplot2 in R.

The resulting PCA projection onto the PC1-PC2 plane is presented in Figure 5, where each data point corresponds to a cluster centroid. Points are color-coded to denote their classification: red for Archaea references, green for Bacteria references, and blue for “Unknown” clusters. The visualization reveals a clear organization among the three classes, visually delineated by three ellipses. These classes occupy distinct spatial regions along the primary axis: the Unknown class is characterized by low PC1 values, the Bacteria reference class spans medium to high PC1 values, and the Archaea reference class exhibits the highest PC1 values, with an overlapping zone situated approximately between PC1 values of 0 and 25. Notably, the first principal component (PC1) accounts for a remarkably high proportion of the explained variance (48.4%), strongly underscoring the statistical significance of this spatial separation. This distinct clustering pattern serves a dual purpose: first, it validates the consistency of our methodology, as the classification coherently aligns with established taxa by grouping reference-annotated clusters together; second, it establishes this analytical framework as a robust tool for exploring bacterial diversity within the 16S rRNA V1-V2 region.

**Figure 5:**
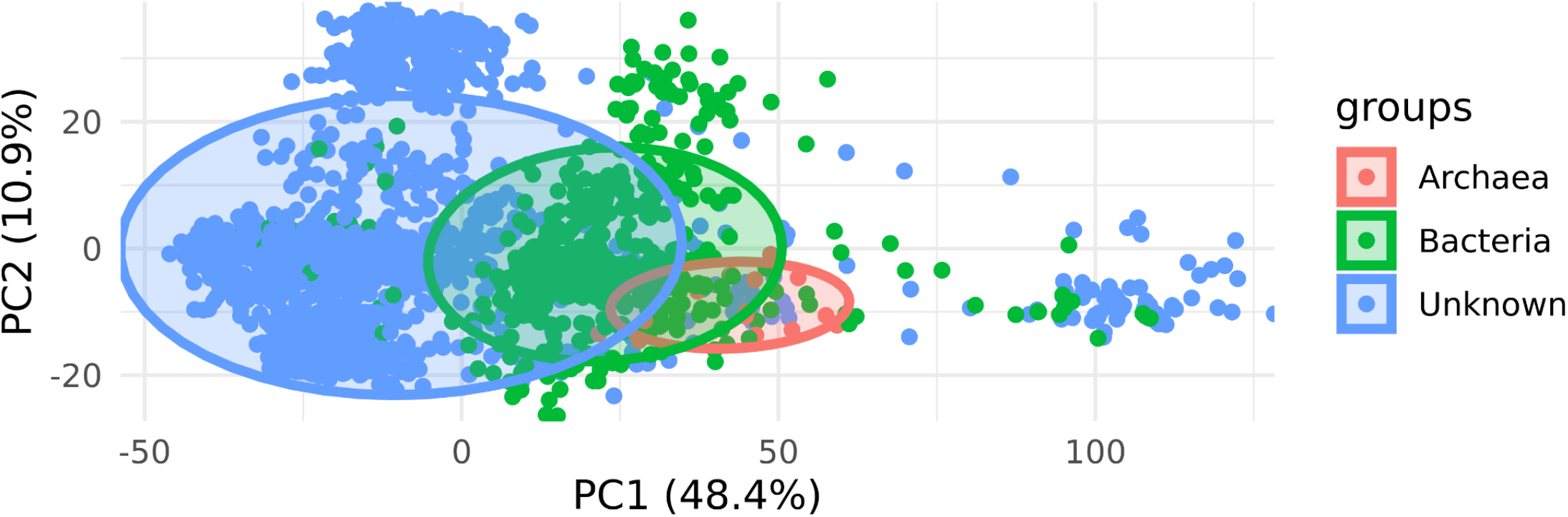
Principal Component Analysis of cluster centroid. Each data point corresponds to a cluster centroid projected onto the PC1-PC2 plane, derived from pairwise alignment distances. Points are color-coded by their classification: Archaea references (red), Bacteria references (green), and Unknown clusters (blue). The visualization demonstrates a clear spatial organization and separation delineated by confidence ellipses along the primary axis (PC1, explaining 48.4% of the variance). Unknown clusters map predominantly to low PC1 values, distinct from the medium-to-high PC1 values of Bacteria and the highest PC1 values of Archaea.

Interestingly, the clusters do not exhibit a uniform distribution across the analyzed sequence space. While the majority of points are concentrated within the central coordinate box [-50:50, -20:20], additional, distinct clouds of points are situated in peripheral regions. Specifically, secondary aggregations are observed at coordinates [-25:10, 20:40] for the Unknown clusters and [25:50, 20:40] for the Reference clusters. Furthermore, an isolated group of points characterized by PC1 values exceeding 75 is located substantially far from the primary point cloud. This distinct outlier group, which warrants further investigation, comprises 58 Unknown clusters and 14 Reference clusters. Taxonomic analysis of these 14 reference clusters revealed the following composition: 6 *Bacillota*, 4 *Pseudomonadota*, 3 *Actinomycetota*, and 1 *Bacteroidota*.

### Classification of samples based on clusters

For every 16S rRNA sample evaluated in this study, all constituent reads were uniquely assigned to the generated clusters, thereby producing a binary vector representation for each sample.

Specifically, each sample is defined by a binary vector with a dimensionality of 43,749 (corresponding to the total number of identified clusters). Within this vector, a value of 1 denotes the assignment of at least one sequence read from the sample to a given cluster, whereas a value of 0 indicates the complete absence of reads mapping to that cluster for the respective sample.

Subsequently, a PCA was conducted to project the samples into the PC1-PC2 coordinate space, facilitating the visual assessment of the distribution across the different sample classes. Figure 6A illustrates the PCA projection of the 734 samples, spanning the three distinct environmental classes detailed in the Materials and Methods section, based exclusively on the presence/absence matrix of the 11,152 annotated clusters. Conversely, Figure 6B presents a parallel PCA executed using the comprehensive presence/absence matrix encompassing all 43,749 clusters.

**Figure 6:**
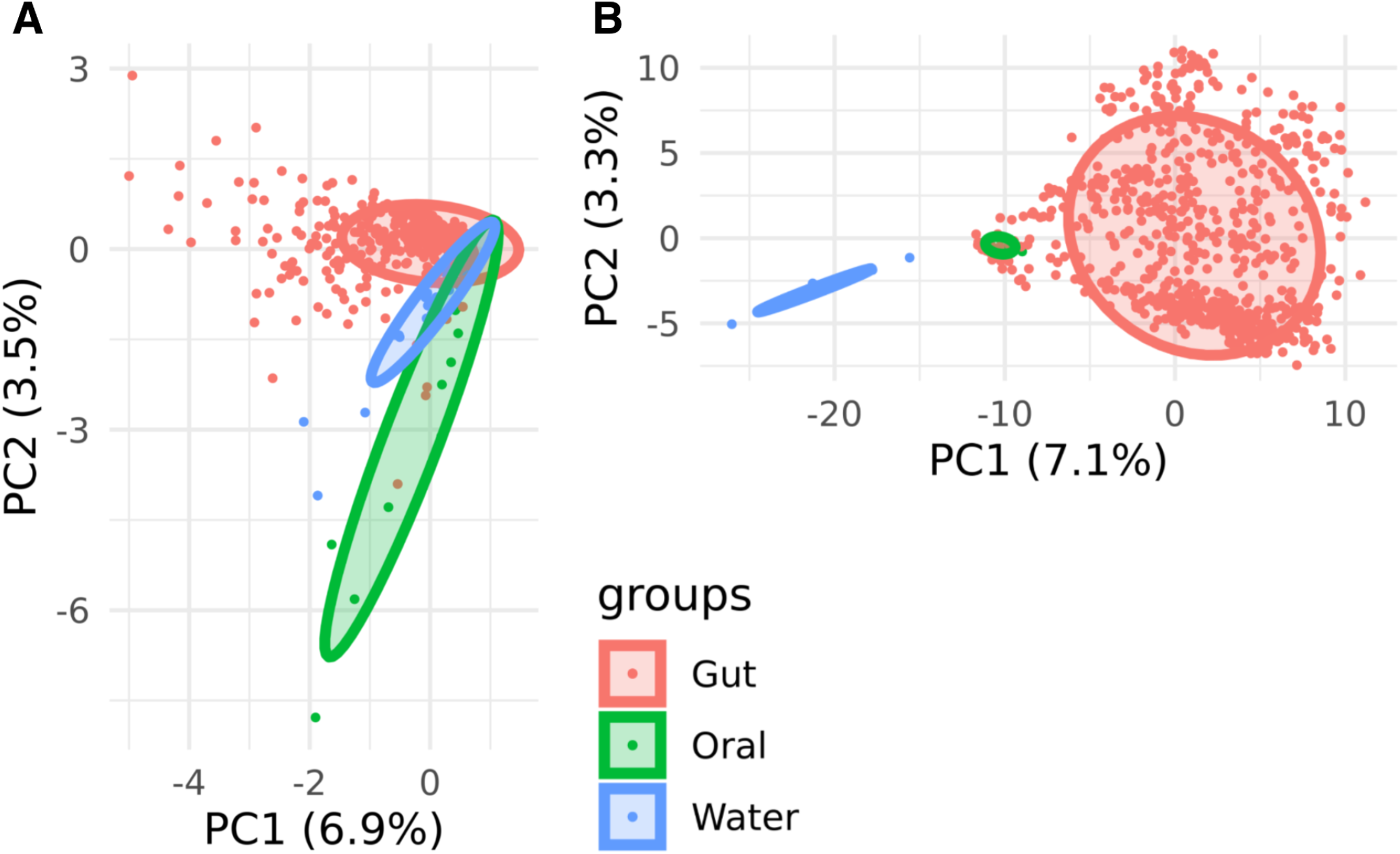
PCA projection of 16S rRNA samples based on cluster occurrence. Samples are projected into the PC1-PC2 coordinate space based on binary presence/absence matrices and color-coded by their environmental origin (Gut, Oral, Water). <u>Panel A</u> – PCA utilizing only the 11,152 annotated (known) reference clusters, which results in partial overlap and ambiguous boundaries between the environmental classes. <u>Panel B</u> – PCA utilizing the comprehensive matrix of all 43,749 clusters, including the newly discovered unknown clusters. The inclusion of the unknown microbial dark matter significantly improves structural resolution, yielding distinctly isolated ecological groups and a biologically consistent representation of the niches.

While the three environmental groups exhibit general spatial separation in both analytical models, Figure 6A demonstrates a partial overlap between the Oral and Water samples. Furthermore, it reveals a central region where all three groups converge, resulting in the intersection of their confidence ellipses. In contrast, the inclusion of all clusters in Figure 6B resolves these ambiguities: the Water samples emerge as a more homogeneous and distinctly isolated group, while the Oral samples are positioned at the periphery of the Gut sample ellipse. This latter spatial arrangement yields a substantially more biologically consistent representation of the relationships among the samples.

Collectively, these findings underscore the pivotal role that uncharacterized bacterial clusters play in structuring and defining metagenomic profiles. Analytical approaches relying solely on previously annotated bacterial clusters may be insufficient for a comprehensive characterization of microbial communities. Conversely, integrating these newly discovered, unknown genera into the analysis affords a significantly more complete, accurate, and representative description of the samples.

### Cluster enrichment

For every cluster, we quantified the number of samples within each environmental group that contained at least one read mapped to that cluster. These raw counts were subsequently normalized into frequencies by dividing them by the total number of samples comprising each respective class. Consistent with the clear spatial separation observed in the PCA projection, the three environmental groups were characterized by distinct cluster occurrence patterns.

An initial analysis focused on isolating highly specific clusters, defined as those appearing exclusively within a single environmental class at a high frequency while remaining completely absent from the others. We identified 152 clusters exhibiting a frequency greater than 0.75 in Water samples, yet entirely absent in both Oral and Gut samples. Notably, only a single cluster within this subset (ID=3166) possessed a known taxonomic annotation, whereas the remaining 151 were classified as unknown. Further stratification of these Water-specific clusters at increasingly stringent frequency thresholds revealed 131 clusters with a frequency exceeding 0.80 (at which point the single annotated cluster, ID=3166, was excluded), 90 clusters at >0.85, 71 at >0.90, 46 at >0.95, and 26 clusters present in 100% of the Water samples (frequency=1.0). All clusters within these subsets remained completely undetected in the Oral and Gut environments.

Conversely, only three clusters, all of which were unknown, exhibited a frequency exceeding 0.75 in the Gut samples while being entirely absent from the Oral and Water classes. Furthermore, no clusters were found to occur with a frequency greater than 0.75 exclusively in the Oral samples. This distribution is biologically plausible, as the Gut and Oral microbiomes are expected to share a closer compositional relationship relative to the Water ecosystem. Additionally, it is important to consider that the statistical representation of the Gut class is inherently influenced by its significantly larger sample size compared to the other environments. Reflecting this shared host-associated overlap, 13 clusters were found at a frequency higher than 0.75 in both Gut and Oral samples while being completely absent from the Water samples.

Finally, to provide a comprehensive visual overview of cluster-sample composition, we isolated the clusters demonstrating a prevalence greater than 0.90 in at least one of the three environmental classes. The abundance profiles of these highly prevalent clusters are visualized as a heatmap in Figure 7, generated using the pheatmap package in R.

**Figure 7:**
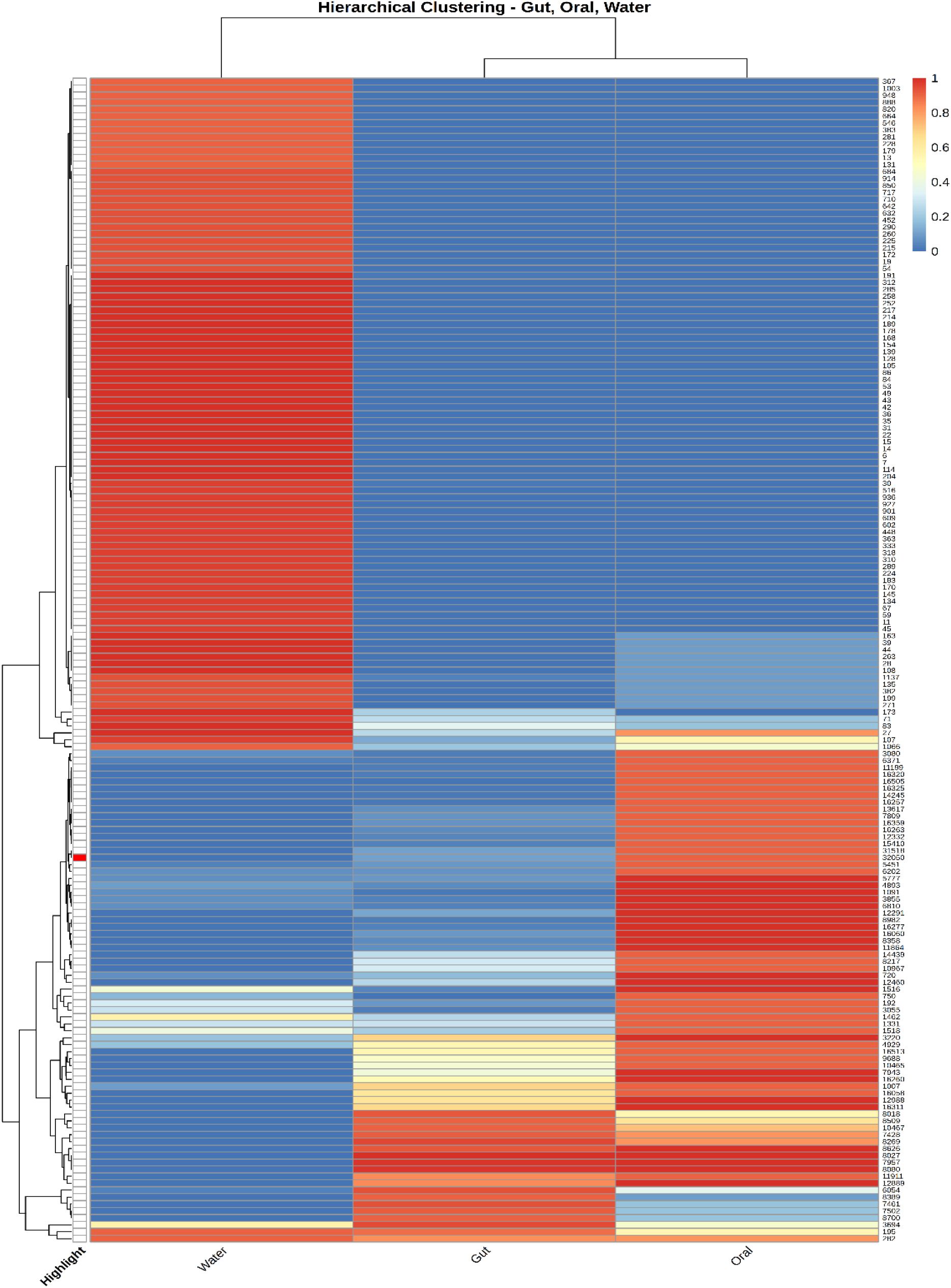
Heatmap of highly prevalent clusters across environmental microbiomes. The heatmap visualizes the normalized occurrence frequencies of specific clusters demonstrating a high prevalence (greater than 0.90) in at least one of the three environmental classes (Water, Gut, Oral). Hierarchical clustering reveals distinct compositional profiles and strong niche specialization, notably highlighting the extensive subset of unannotated clusters that are exclusively and highly enriched in the Water ecosystem compared to host-associated environments.

## DISCUSSION AND CONCLUSIONS

The exponential accumulation of 16S rRNA gene sequencing data in public repositories represents a historic opportunity to catalog global microbial diversity. However, the computational limitations of traditional, all-vs-all clustering algorithms have forced researchers to rely on static, pre-computed reference databases, effectively blinding them to the vast reservoir of uncharacterized microbes. In this study, we introduced Meta16S, a highly scalable, reproducible, and dynamically expanding framework designed to overcome these bottlenecks. By shifting the paradigm from static clustering to an incremental approach, Meta16S enables the continuous *de novo* discovery and cataloging of “microbial dark matter” from unending streams of data.

The application of our pipeline to a diverse corpus of 734 human and environmental samples yielded striking results that underscore the severe limitations of current reference-based profiling. Our initial reference database, rigorously curated from over 56,000 reference genomes, yielded 12,150 distinct, annotated known clusters at the 95% identity threshold. However, upon incrementally processing the 16S rRNA samples, the database expanded massively, resulting in a final catalog of 43,749 clusters. This indicated that 72.23% of the OTUs identified in this relatively modest dataset represent unknown taxa. The sheer volume of this unclassified diversity confirms that existing genomic databases capture only a fraction of the true microbial world. Meta16S provides the computational infrastructure necessary to capture and formalize this missing diversity into stable, reusable identifiers.

Beyond simply quantifying this unknown diversity, our sequence-level PCA demonstrated that these novel clusters occupy distinct regions of the 16S V1-V2 sequence space. The clear separation between known Bacteria, known Archaea, and the newly discovered unknown clusters suggests that much of this microbial dark matter is not merely composed of minor variants of known species, but rather represents deeply branching phylogenetic lineages. Furthermore, the identification of a subset of clusters, including 58 highly divergent unknown OTUs, located far from the primary sequence clouds hints at the discovery of highly novel extremophiles or deeply isolated evolutionary branches that warrant targeted biological characterization in future studies.

Perhaps the most compelling finding of this study is the profound ecological significance of these unknown clusters. This was clearly demonstrated when classifying the 16S rRNA samples themselves. When mapping samples exclusively against the annotated (known) database, the resulting PCA showed ambiguous boundaries, particularly between oral and environmental water samples. However, when the unknown clusters were introduced into the presence/absence matrix, the resolution improved dramatically. The water samples collapsed into a distinct, homogeneous cluster entirely separate from the host-associated samples, while the unknown microbes are not simply sequencing noise or biological artifacts. They carry highly specific, environment-defining ecological signals. Ignoring them inherently compromises the accuracy of community profiling.

This niche specialization was further validated by our cluster enrichment analysis. We observed a striking number of unknown clusters that were highly restricted to specific ecosystems. For instance, the identification of 151 unknown clusters that appear exclusively and with high frequency (>75%) in water samples highlights a known bias in current reference databases, which are heavily skewed toward human-associated and easily culturable pathogens [48–50]. By capturing these highly prevalent but uncultivated environmental microbes, Meta16S bridges the gap between clinical microbiology and broad environmental ecology, providing researchers with the markers needed to track taxa across diverse biomes.

While Meta16S provides a powerful step forward, we acknowledge certain limitations inherent to amplicon sequencing. Our current implementation relies on the V1-V2 hypervariable region, which, while highly robust for species and genus-level discrimination, is bounded by the inherent limitations of a single marker gene. Furthermore, our use of a 95% identity threshold serves as a necessary heuristic proxy for genus-level clustering, though biological boundaries are rarely strictly defined by fixed distance metrics. Nevertheless, the modular nature of the Snakemake workflow ensures that future iterations can easily be adapted to different hypervariable regions, full-length 16S sequences generated by long-read technologies, or alternative clustering thresholds.

In conclusion, the Meta16S framework represents a fundamental shift in how large-scale microbiome data is processed and interpreted. It transitions the field away from siloed, static analyses and toward the creation of a continuously evolving, global census of microbial life. By proving that unknown sequence clusters drive the structural separation of ecological niches, we highlight the critical need to include microbial dark matter in routine taxonomic profiling. Looking forward, the scalability of this pipeline facilitates large-scale analysis of public repositories like the Sequence Read Archive, supporting the ongoing development of more comprehensive microbial reference databases.

## ADDITIONAL INFORMATION

### Availability

All code and data generated for this study are publicly available to ensure full reproducibility and to facilitate reuse by the community.

The complete Snakemake workflow, including all scripts and the Conda environment file, is available on GitHub under the MIT license at https://github.com/cumbof/Meta16S, while the Meta16S databases are maintained on Zenodo at https://doi.org/10.5281/zenodo.20087185.

Meta16S requires Snakemake as a workflow management engine, VSEARCH v2.30.0 for the filtering, dereplication, chimera removal, and the incremental clustering of 16S sequences, barrnap v0.9 for the identification of 16S regions from Bacteria and Archaeal reference genomes, and ncbitax2lin v2.4.1 for the retrieval of taxonomic information of reference genomes from NCBI GenBank. These requirements are all open-source and installable via Conda from the Bioconda channel (*conda install -c bioconda snakemake vsearch barrnap ncbitax2lin*).

While the processed cluster databases and workflow code are publicly accessible, the raw 16S rRNA amplicon sequencing samples supporting the findings of this study are available from the authors upon request.

### Author Contributions

FC, DS, and VRS conceived the research; FC and DS implemented the workflow, and curated the Meta16S database; FC, DS, GF, DB, FV, and VRS contributed to the analysis and discussion of the Meta16S clustering and classification results, wrote the manuscript, and agreed with its final version.

### Conflict of Interest

No conflicts to disclose.

## Acknowledgments

Finally, we would like to acknowledge the use of a Large Language Model (Google’s Gemini 3 Pro) in refining the clarity and readability of this manuscript. The AI assistance was primarily used for tasks such as sentence restructuring, word choice suggestions, and identifying potentially unclear phrasing. We emphasize that the AI was used solely for language enhancement and did not contribute to the generation of research ideas, data analysis, or the interpretation of results. All conclusions drawn and insights presented in this manuscript are solely the product of the authors’ own analysis and expertise.

## Notes

https://github.com/cumbof/Meta16S

### Competing Interest Statement

The authors have declared no competing interest.

